# Dating the origin and spread of specialization on human hosts in *Aedes aegypti* mosquitoes

**DOI:** 10.1101/2022.09.09.507331

**Authors:** Noah H. Rose, Athanase Badolo, Massamba Sylla, Jewelna Akorli, Sampson Otoo, Andrea Gloria-Soria, Jeffrey R. Powell, Bradley J. White, Jacob E. Crawford, Carolyn S. McBride

## Abstract

The globally invasive mosquito subspecies *Aedes aegypti aegypti* is a highly effective vector of human arboviruses because it specializes in biting humans and breeding in human habitats. Recent work suggests that specialization first arose as an adaptation to long, hot dry seasons in the West African Sahel, where *Ae. aegypti* is forced to rely on human-stored water for breeding. However, rainfall patterns in this region have changed dramatically over the past 10-20 thousand years, and we do not yet know exactly when specialization occurred. Here we use whole-genome cross-coalescent analysis to date the emergence of human specialist populations in the Sahel and thus further probe the climate hypothesis. Importantly, we take advantage of the known migration of human-specialist populations out of Africa during the Atlantic Slave Trade to calibrate the coalescent clock and thus obtain a more precise estimate of the older evolutionary event than would otherwise be possible. We find that human-specialist mosquitoes diverged rapidly from ecological generalists approximately 5,000 years ago, which corresponds to the end of the African Humid Period—a time when the Sahara dried and water stored by humans became a uniquely stable, aquatic niche in the Sahel. We also use population genomic analyses to date a previously observed influx of human-specialist alleles into major West African cities, where mosquitoes tend to be more attracted to humans than in nearby rural populations regardless of climate. In this case, the characteristic length of tracts of human-specialist ancestry present on a generalist genetic background in Kumasi, Ghana and Ouagadougou, Burkina Faso suggests the change in behavior occurred during rapid urbanization over the last 20-40 years. Taken together, we show that the timing and ecological context of two previously observed shifts towards human biting in *Ae. aegypti* differ; climate was likely the original driver, but urbanization has become increasingly important in recent decades. Understanding the changing relationship between mosquitoes and humans over time is critical for predicting and managing burdens of mosquito-borne disease.

## Introduction

The mosquito *Aedes aegypti* is thought to have originated as an ecological opportunist in forested areas of sub-Saharan Africa, and extant African populations of the subspecies *Ae. aegypti formosus* are relative generalists (Figure 1A). They breed in a wide variety of habitats and opportunistically bite a wide variety of human and non-human animals (*1*–*4*). However, at some point in the past, a few populations in West Africa evolved to specialize in living with and biting humans, giving rise to a human-adapted subspecies *Ae. aegypti aegypti* that has since spread around the global tropics (Figure 1A) and become the world’s most effective vector of the viruses responsible for yellow fever, dengue, Zika, and chikungunya (*1, 5, 6*, but see *7*).

**Figure 1.**
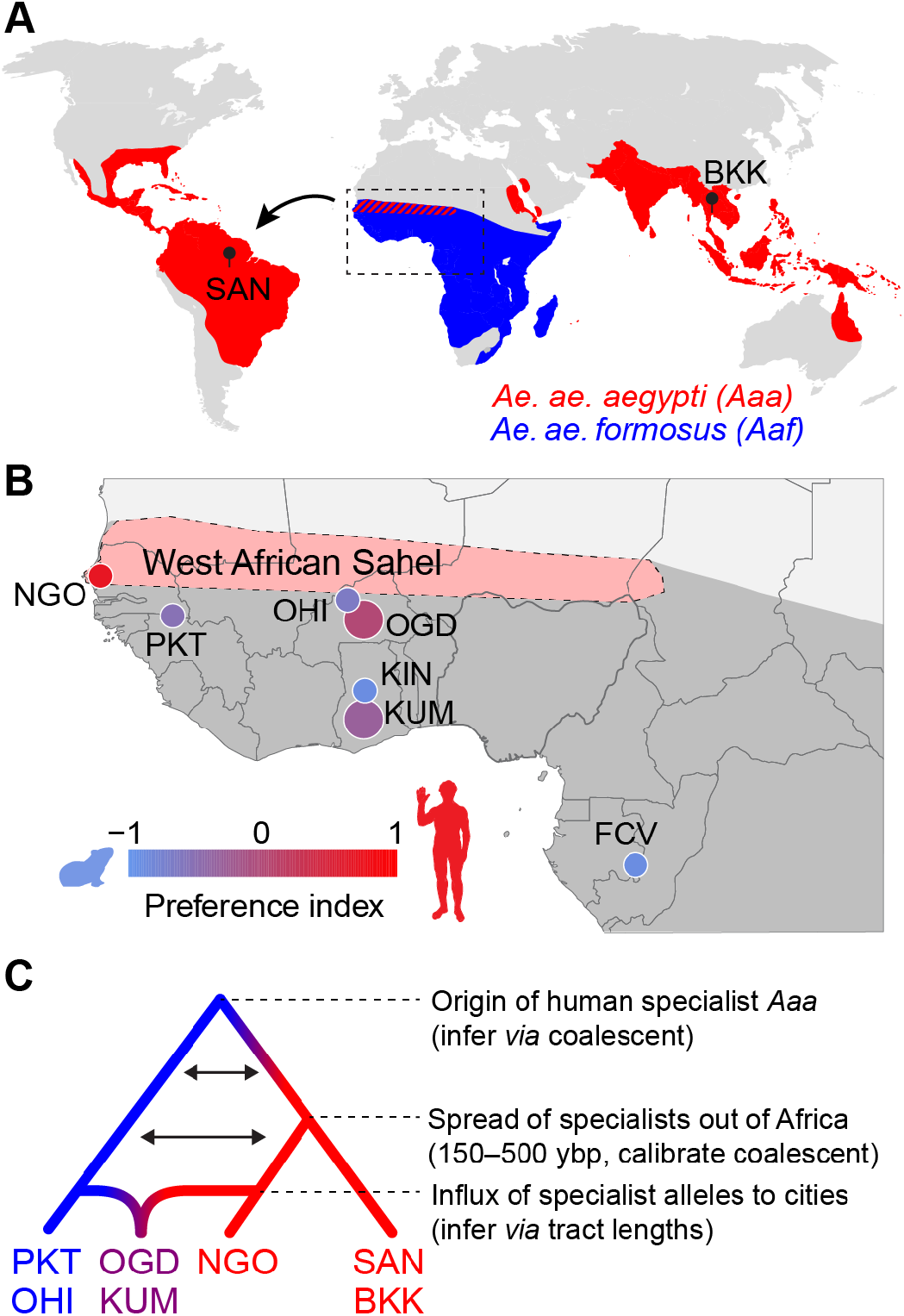
Dating the origin and spread of the human-specialist form of *Aedes aegypti*. **A**. The human-specialist form of *Aedes aegypti* (*Aaa*) is thought to have originated in West Africa, before invading the Americas in association with the Atlantic Slave Trade and subsequently spreading across the global tropics. **B**. Present-day *Ae. aegypti* populations in West Africa vary in their preference for human hosts. Large cities (e.g. KUM, OGD) are shown as larger circles, while small towns are shown as smaller circles. Map region corresponds to the inset in panel A. Pink shading marks the intensely seasonal Sahel region, where human specialists likely originated. Behavioral data taken from Rose et al. 2020. **C**. The timing of major events in the evolutionary history of human specialist populations (red lineages) can be inferred using population genomic approaches, such as coalescent analysis (for older events) and tract length analysis (for more recent events). SAN, Santarem, Brazil; BKK, Bangkok, Thailand; NGO, Ngoye, Senegal; PKT, PK10, Senegal; OHI, Ouahigouya, Burkina Faso; OGD, Ouagadougou, Burkina Faso; KUM, Kumasi, Ghana; KIN, Kintampo, Ghana; FCV, Franceville, Gabon.

Understanding the ecological factors that drive mosquitoes like *Ae. aegypti* to adapt to human hosts and habitats is important because it can help us predict and respond to future burdens of mosquito-borne disease on a planet whose climate and landscape is being transformed by human activity. Recent work suggests that human specialization in *Ae. aegypti* originally evolved as an adaptation to the extremely long, hot dry seasons of the Sahel region of West Africa (Figure 1B). *Aedes* mosquitoes lay their eggs just above the water line in small, contained bodies of water, and immature stages require water for development. The scarcity of natural tree-hole and rock-pool aquatic habitat for up to 9 months of the year in the Sahel likely forced *Ae. aegypti* to rely on human-stored water for survival, with adaptations to breeding in artificial containers (*6, 8*–*11*) and biting humans (*1, 2, 12*) soon following. However, neither highly seasonal precipitation patterns nor human-derived pots of water were present in the Sahel until relatively recently in evolutionary time. Settled human societies practicing water storage in clay vessels likely developed within the last 10,000 years in sub-Saharan Africa (*13, 14*). Meanwhile, the Sahara Desert and associated Sahelian biome have only existed in their present form for the past 5,000 years; the region was a vast savannah during the African Humid Period from 15,000 to 5,000 years ago. If *Ae. aegypti* evolved to specialize in biting humans as a by-product of breeding among humans during the intense dry seasons of the West African Sahel (hereafter “Sahel”), then the shift should have occurred no more than 5,000 years ago.

The idea that *Ae. aegypti aegypti* first emerged when all elements of its current niche were finally present approximately 5,000 years ago has been suggested multiple times in the vector biology literature (*e.g. 6, 9*), but has never been comprehensively tested with modern genomic data. A site-frequency-spectrum-(SFS-) based analysis of exome sequence data estimated that neighboring generalist and human-specialist populations in West Africa diverged about 15,000 years ago (*15*), but this estimate was sensitive to model specification and parameterization. Conversely, a molecular clock-based phylogenetic analysis found that all African populations of *Ae. aegypti* have a common ancestor approximately 17,000 to 25,000 years ago, but did not precisely date the subsequent split between human-specialist and generalist subspecies within Africa (*16*).

In contrast to the uncertainty surrounding exactly when (and why) specialization occurred (Figure 1C), the historical forces that led to the spread of *Ae. aegypti aegypti* out of Africa are relatively well understood. Human-specialist *Ae. aegypti* is thought to have first migrated to the Americas on ships during the Atlantic Slave Trade approximately 500 years ago (Figure 1C). Its arrival there coincided with the first recorded outbreaks of yellow fever in the New World in the 1600s (*6*). Subsequent outbreaks of yellow fever exacted an enormous toll in the following centuries, especially among immunologically naive European and Indigenous populations, shaping not just health outcomes, but also the political and economic history of the Americas (*17*). Similarly, *Ae. aegypti*’s invasion of Asia and Oceania was followed by outbreaks of dengue, chikungunya, and later Zika, although yellow fever remains absent in these areas (*5, 6*). Today, invasive populations of *Ae. aegypti aegypti* in the Americas and Asia are almost all closely related to each other and distinct from their ancestors in Africa (*6, 18*).

A third and putatively much more recent event in the evolutionary history of human-specialist *Ae. aegypti* involves a shift towards human-biting in growing African cities that would otherwise be expected to harbor generalist populations. Regardless of local climate, urban *Ae. aegypti* mosquitoes in places like Kumasi, Ghana and Ouagadougou, Burkina Faso are consistently more responsive to humans than those from nearby rural areas (Figure 1B, compare KUM to KIN and OGD to OHI) (*2*). This shift is associated with the same underlying ancestry component that defines human specialists in the Sahel and outside Africa, suggesting that it results from an influx of human-specialist alleles into urban areas rather than independent evolution of human-biting (Figure 1C) (*2*). Although the exact timing of the observed influx in places like Kumasi and Ouagadougou remains to be tested, it likely occurred recently; the cities involved have only reached the high levels of human population density associated with greater mosquito preference for humans within the past few decades (*19*). Urban populations that lack substantial admixture from this human-specialist ancestry component have also been described and posited to represent genetically independent adaptation to human environments (*20*). However, when their behavior has been tested, these populations do not show increased preference for human hosts (*2*). This does not preclude other forms of adaptation to human habitats.

Here, we take advantage of existing population genomic data from diverse African and non-African populations (*2*) to infer the timing of two key events in the evolutionary history of the human-specialist form of *Ae. aegypti* (Figure 1C) and thereby more rigorously test key hypotheses concerning their underlying ecological drivers. First we address the timing of initial emergence of specialist populations in the Sahel *via* cross-coalescent analysis—a methodological framework developed for studying human evolution that has rarely been applied in other species. Importantly, we take advantage of the known migration of specialist *Ae. aegypti* out of Africa 150-500 years ago to calibrate the coalescent clock and thus obtain a much more precise estimate than would otherwise be possible. Second, we use ancestry tract length analysis to infer the timing of the recent shift towards human-biting in large West African cities. We find that human specialists originated with the emergence of the modern Sahelian climate approximately 5,000 years ago, while the influx of human-specialist ancestry into urban areas coincides with rapid urbanization over the past 20-40 years.

## Results

### Dating the origin of human specialists within *Ae. aegypti*

One way to date the evolution of specialization on humans in *Ae. aegypti* is to examine the time course of divergence between specialist and generalist populations using cross-coalescent analysis. Coalescent approaches model relationships between haplotypes at a locus going backwards in time, using the steady accumulation of mutations to estimate when haplotypes merge into a single ancestral sequence (*21*). Using the Multiple Sequentially Markovian Coalescent (MSMC), one can characterize the distribution of times to coalescence across genome-wide loci (*21*). Patterns of relative cross-coalescence are particularly informative. This is the rate of coalescence of haplotypes from different populations relative to the rate of coalescence of haplotypes from the same population. It should start close to zero in the recent past when two populations are isolated, but eventually plateau at one when, going back in time, they have merged into a single ancestral population. Given a known mutation rate and generation time, patterns of relative cross coalescence can be used to infer the timing of splits and subsequent gene flow between populations (*21*).

One hurdle associated with effective application of cross-coalescent analysis to real world genomic data is the need for phased genomes. To meet this challenge, we leveraged statistical approaches and recently published short-read sequence data for 389 unrelated *Ae. aegypti* individuals (*2*) to generate a large phasing panel for this species. First, we used *HAPCUT2* to pre-phase individuals (*22*). In this step, we used linkage information present in short reads and read pairs to locally phase adjacent variants; because nucleotide diversity in *Ae. aegypti* is high (about 2% (*2*)), multiple variants are often present within a single sequencing fragment, making pre-phasing highly effective in this species. Typical local haplotypes, or “phase sets,” ranged from hundreds of bases to a few kilobases. We then used *SHAPEIT4*.*2* to assemble local haplotypes into chromosome-level haplotypes, using statistical linkage patterns present across our panel of 389 individuals (*23*).

A second hurdle associated with coalescent analysis is the need for a reliable scaling factor that can be used to translate coalescent time into real time. This parameter depends on the *de novo* mutation rate and generation time, neither of which is precisely known for *Ae. aegypti*. The mutation rate is likely on the order of 3×10^−9^ per generation given estimates from other insects (*Drosophila melanogaster:* 2.8×10^−9^, *Bombus terrestris:* 3.6×10^−9^, *Heliconius melpomene:* 2.9×10^−9^) (*24*–*26*), and most populations of *Ae. aegypti* probably go through ∼15 generations per year (*15, 18, 27, 28*). However, there is substantial uncertainty in both numbers (10 or 12 generations per year are also common estimates, for example (*18*)), and any estimate of the timing of specialization derived from these values would be extremely rough.

To overcome this problem, we decided to leverage the known timing of the spread of human-specialist *Ae. aegypti* out of Africa during the Atlantic Slave Trade to ground-truth and calibrate our coalescent scaling factor. If the species did indeed escape Africa at this time, coalescent analysis of the relationship between human specialist populations within and outside Africa should show a single strong pulse of migration that matches the timing of the Atlantic Slave Trade for plausible values of the mutation rate and generation time. If we then assume these values are constant through evolutionary time, we can use them in subsequent analyses of the older split between specialists and generalists.

We used *MSMC2* and *MSMC-IM* to fit an isolation-with-migration model of the relationship between a human-specialist population from the Sahel (Ngoye, Senegal [NGO], Figure 1B) and a human-specialist population from the invasive range (Santarem, Brazil [SAN], Figure 1A) (*21*). As expected, this analysis revealed a single strong pulse of migration (Figure 2A, inset), which we presume corresponds to the Atlantic Slave Trade. NGO has experienced substantial admixture from nearby generalist populations, and is almost certainly not the sole origin point of invasive populations. For example, populations in Luanda, Angola are more closely related to invasive populations in the Americas (*29*) (see Discussion). Nevertheless, NGO provides the best available proxy for human-specialist populations in their native habitat, and the observed migration signal should reflect the time course of their spread to the Americas.

**Figure 2.**
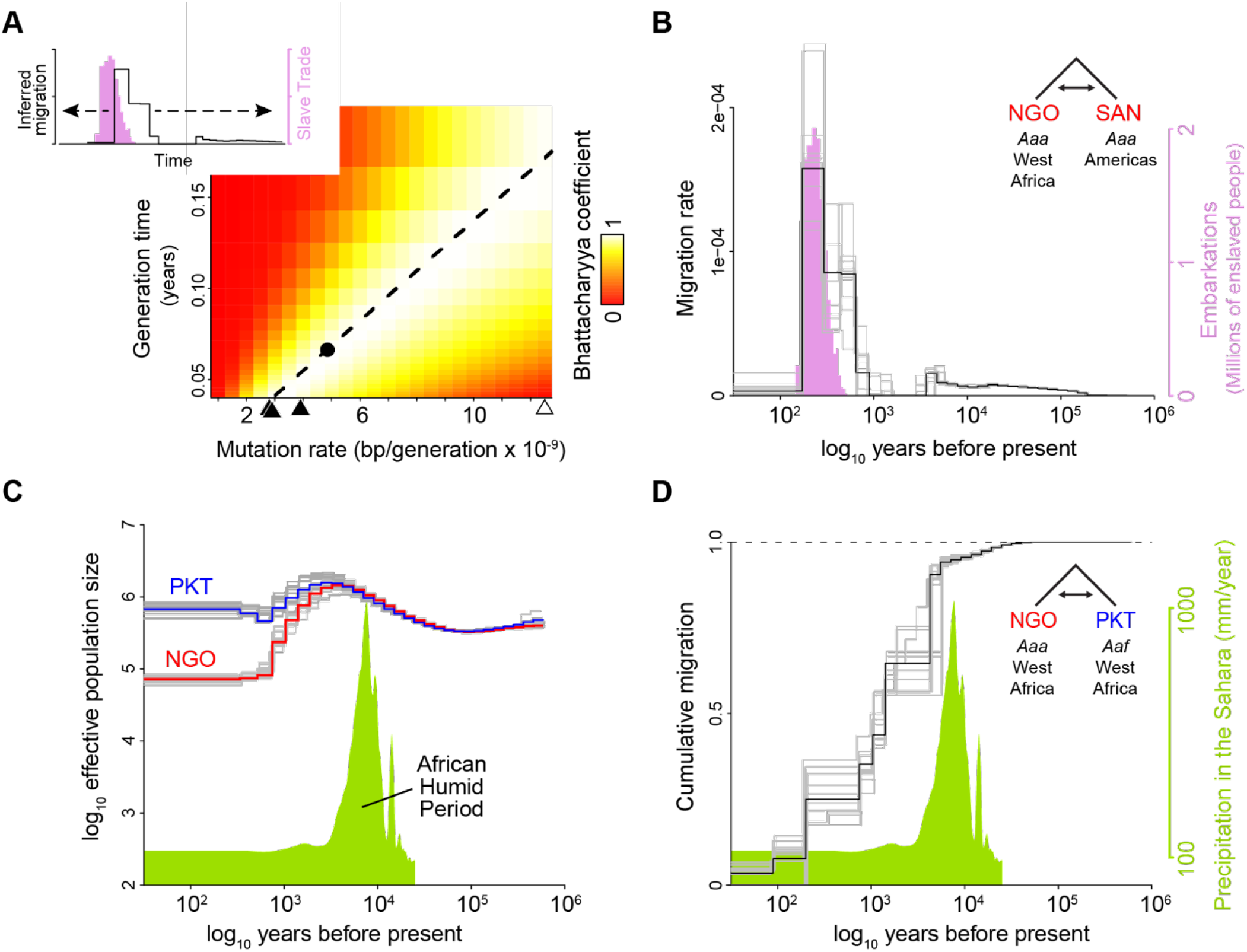
A calibrated coalescent scaling factor for *Ae. aegypti* suggests rapid evolution of human specialists at the end of the African Humid Period. **A**–**B**. Calibration of a coalescent scaling factor for *Ae. aegypti*. **A**, The Bhattacharyya coeficient reveals the extent of overlap between the timing of the Atlantic Slave Trade (based on historical records (*30*), pink distribution in inset) and inferred migration of *Aaa* out of Africa (based on cross-coalescent analysis, black line in inset) for given combinations of the mutation rate (μ) and generation time (g). A value of 1 indicates perfect correspondence through time and 0 indicates no overlap. Estimated mutation rates for three other insects (see Results) and *Homo sapiens* are marked along x-axis with closed and open triangles, respectively. y-axis ranges from the shortest to longest reasonable generation times for *Ae. aegypti*. The dashed line indicates the μ/g ratio that provides the strongest match between genomic and historical data. The dot along this line marks the calibration used in the other panels (g=0.067, μ=4.85×10^−9^). **B**. The inferred timecourse of migration of *Aaa* from West Africa to the Americas shown alongside the historical record of the Atlantic Slave Trade that was used for calibration. **C**–**D**. Cross-coalescent analysis of the timing of human specialization. **C**. Inferred effective population sizes for West African *Aaa* (NGO) and *Aaf* (PKT) superimposed on Saharan rainfall data (inferred from Atlantic marine sediments (*29*)). **D**. Estimated cumulative migration between *Aaa* (NGO) and *Aaf* (PKT) is expected to plateau at 1 going backwards in time, when populations have merged into a single ancestral population. Populations diverged rapidly at the end of the African Humid Period.

We next obtained publicly available data on the number of enslaved people trafficked across the Atlantic in 25-year intervals between 1500 and 1875 (*28*) and used the Bhattacharyya Coefficient to assess the degree of correspondence between this historical record and *MSMC*-inferred mosquito migration for given combinations of the mutation rate and generation time (Figure 2A). The Bhattacharyya Coefficient should take a value of zero in the case of non-overlapping distributions (red areas of plot) and one in the case of perfect overlap (white areas). The fact that we see good overlap between the two distributions (yellow–white color) across a wide range of reasonable mutation rates and generation times for *Ae. aegypti* is consistent with our understanding of the species’ recent history and supports our approach. For example, if we take the common literature value of 15 generations per year (0.067 years per generation) (*15, 18*), the *de novo* mutation rate that maximizes correspondence between the two datasets is 4.85×10^−9^ (black dot in Figure 2A, used in Figure 2B), which is on the order of values documented in other insects. We chose to carry forward this calibrated scaling factor into subsequent analyses. Note that using Bangkok, Thailand (BKK, Figure 1A) as our reference invasive population instead of Santarem, Brazil produced a similar pattern, although the peak of the migration pulse was slightly older (Figure S1L *versus* P). In both cases, time-calibrated demographic histories showed a bottleneck in invasive populations about 500 years ago (Figure S1A,B).

We next sought to examine the historical relationship between human-specialist and generalist populations of *Ae. aegypti* within Africa using our newly calibrated scaling factor. We again used *MSMC2* and *MSMC-IM* to characterize patterns of cross-coalescence, but this time contrasting human-specialists in Ngoye, Senegal (NGO, Figure 1B) with nearby generalists from PK10, Senegal (PKT, Figure 1B). We infer that these populations had similar effective population sizes ∼10,000 years ago, which gradually expanded before diverging about 5,000 years ago (Figure 2C). Isolation-with-migration analysis also suggested that these populations abruptly diverged at about 5,000 years before present. This can be seen in the cumulative migration signal, which should reach one when (going back in time) populations have merged into a single ancestral population. Cumulative migration begins dropping slowly 15-30 thousand years ago, suggesting a small amount of genetic structure in ancient times, but then drops precipitously 3-5 thousand years ago (Figure 2D). This rapid divergence shows a striking correspondence to the end of the African Humid Period, as evidenced by the record of leaf wax isotopes present in marine sediments from the Atlantic Ocean floor (Figure 2D) (*31*). Analyses using an alternative generalist population (OHI, Figure 1B) yielded very similar results (Figure S1).

### Dating the origin of increased human specialization in urban populations

Within Africa, human-specialist populations of *Ae. aegypti* are largely restricted to the unusual Sahel climatic zone (Figure 1A–B). However, even in places where most mosquitoes are generalists, urban populations tend to be more responsive to human hosts than their rural counterparts (*2*). This increased willingness to bite humans is strongly correlated with levels of admixture from a shared human-specialist ancestry component found across locations (*2*), suggesting that this behavioral shift is the result of a recent influx of human specialist alleles into rapidly growing cities. Before estimating the timing of this shift, we used the *f3* statistic to confirm that *Ae. aegypti* populations in the large cities of Kumasi, Ghana (KUM) and Ouagadougou, Burkina Faso (OGD), could be accurately modeled as mixtures of human-specialist and generalist populations (*32*). This statistic tests whether allele frequencies in a focal population are consistently intermediate between frequencies in two candidate source populations. If so, the genome-wide average *f3* should be negative, while a positive value provides evidence neither for nor against admixture. We used three different populations as candidate sources of human specialist ancestry (West African NGO, American SAN, or Asian BKK, Figure 1A,B) and two different populations as candidate sources of generalist ancestry (West African KIN or OHI, Figure 1B). The resulting f3 values were substantially and significantly negative for both cities in almost all cases (Figure 3A,B). However, the use of NGO as a proxy for the human specialist source resulted in only modestly negative values for KUM and positive values for OGD (Figure 3A,B). This ambiguous signal likely reflects the complex history of NGO, which is itself admixed despite its human-specialist ecology. The non-African specialists, in contrast, are geographically isolated from African generalists. They may therefore be a better proxy for an unadmixed human specialist source even if the specialist ancestry present in KUM and OGD came from within Africa. Denser sampling of human specialists both within and outside Africa will be necessary to come to a definitive conclusion about the true sources of admixture in these large cities.

**Figure 3.**
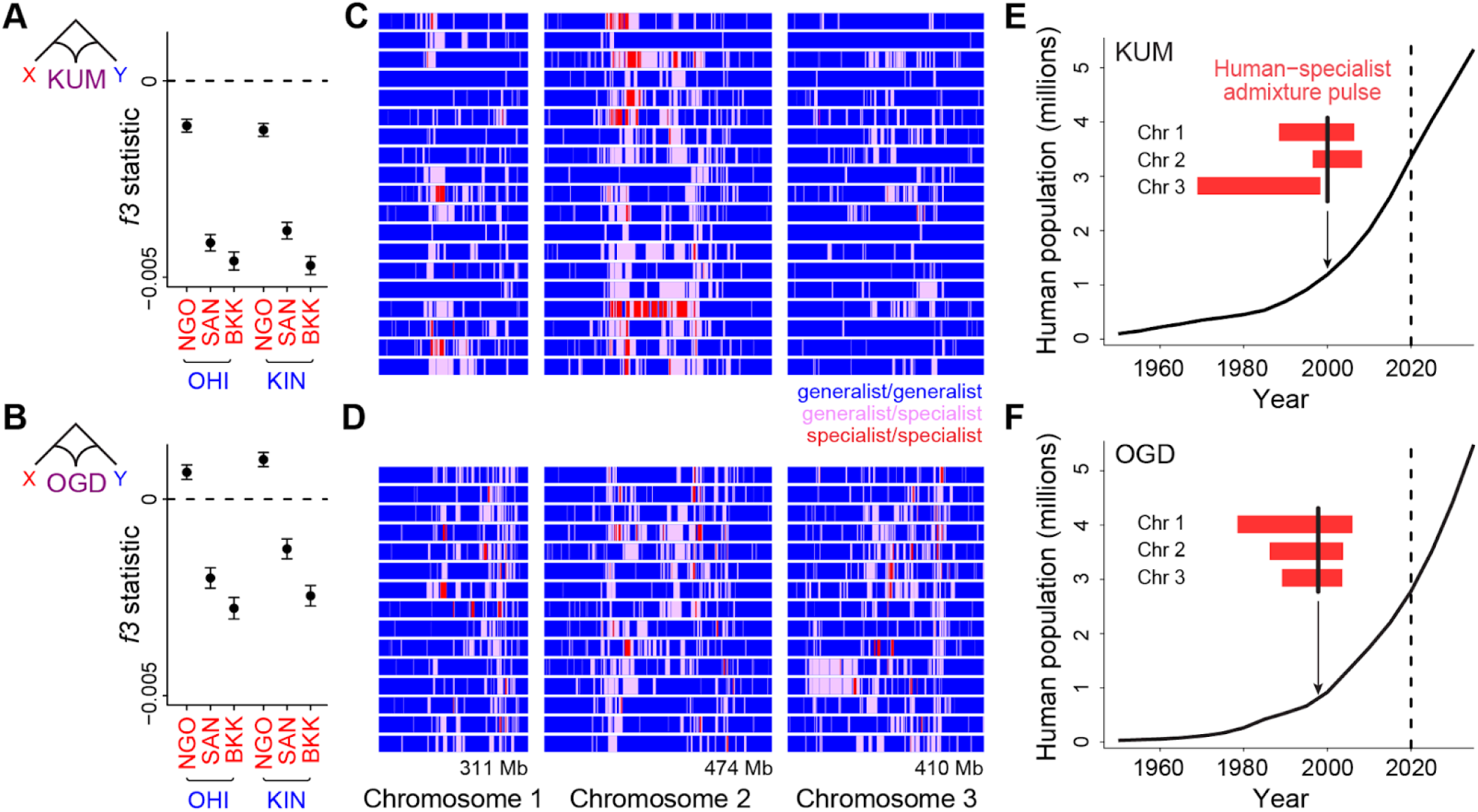
Long tracts of human-specialist ancestry in rapidly growing cities suggest a recent influx associated with modern urbanization. **A**–**B**. *f3* analysis confirming that Kumasi, Ghana (**A**, KUM) and Ouagadougou, Burkina Faso (**B**, OGD) can be modeled as a product of admixture between generalist and human-specialist *Ae. aegypti* populations. Negative values provide evidence of admixture. Error bars show 95% jackknife confidence intervals. **C**–**D**. Distribution of tracts of human-specialist ancestry (purple, heterozygous; red, homozygous) across 19 unrelated KUM genomes (**C**) and 15 unrelated OGD genomes (**D**). **E**–**F**. The inferred timing of human-specialist admixture (red bars) overlaid on publicly available human population growth data and near-future growth projections for KUM (**E**) and OGD (**F**). Red bars correspond to 95% confidence intervals for each chromosome, and thick black lines mark the mean for all three chromosomes.

To test whether the timing of admixture in KUM and OGD matches the timing of the rapid growth of human populations in each location, we turned to ancestry tract length analysis. When human-specialist migrants first come into a generalist population, their offspring will have long tracts of human-specialist ancestry in their genomes (*33*). Over time recombination should break these tracts into smaller and smaller pieces, making tract length an inverse correlate of the amount of time that has passed since the influx of human-specialist ancestry occurred (T3, Figure 1C) (*34*). These patterns can be used to investigate relatively recent gene flow, whereas coalescent approaches are informed by mutations that take thousands of generations to accumulate. We conducted ancestry tract analysis in KUM and OGD, as implemented in *AncestryHMM* (*34*), using as source populations the human specialists and generalists that gave the strongest signals of admixture in our f3 tests: BKK and OHI.

We found long tracts of human-specialist ancestry in KUM and OGD, indicative of a relatively recent influx of human-specialist alleles (Figure 1C,D). Using the recombination map from the *Ae. aegypti* L5 assembly, and masking regions previously identified as under divergent selection between human-specialists and generalists (see Methods) (*2, 35*), we estimated the number of generations of recombination since admixture. In both cities, admixture appears to have been recent, occurring within the last 20-40 years, coinciding with rapid human population growth in both cities (Figure 3E,F). The distribution of human-specialist tracts was not entirely uniform; instead they were more concentrated in a few key regions (Figure 3C,D). This signal could not be explained by failure to detect admixture in specific regions as simulations confirmed that we had statistical power to identify mixing across the whole genome (Figure S2). Moreover, the presence of human-specialist ancestry was more spatially correlated across the genomes of individuals from the two cities than would be expected due to random chance (permutation P<0.001, Figure S3). This signal could reflect parallel patterns of selection for human-specialist ancestry in chromosomal regions that are important for urban adaptation. Overall, our results are consistent with a recent influx of human-specialist ancestry into the rapidly growing cities of Kumasi and Ouagadougou.

## Discussion

Taken together, our results suggest that the ecology of *Ae. aegypti* has been in flux across human history (Figure 4). We previously found that *Ae. aegypti* populations in their native range in sub-Saharan Africa were most specialized on human hosts in the West African Sahel, where they depend on human water storage for breeding across the long dry season (*2*). However, urban populations were also more attracted to human hosts than nearby rural populations regardless of climate (*2*). Here, we used cross-coalescent and ancestry tract analyses of genomic data to determine when these ecological shifts occurred and thus gain additional insight into their underlying drivers. According to our cross-coalescent estimates, human-specialist and generalist populations of *Ae. aegypti* diverged rapidly at the end of the African Humid Period, about 5,000 years ago. In contrast, our ancestry tract analyses suggest that the urbanization effects are much more recent, playing out over the last 20-40 years.

**Figure 4.**
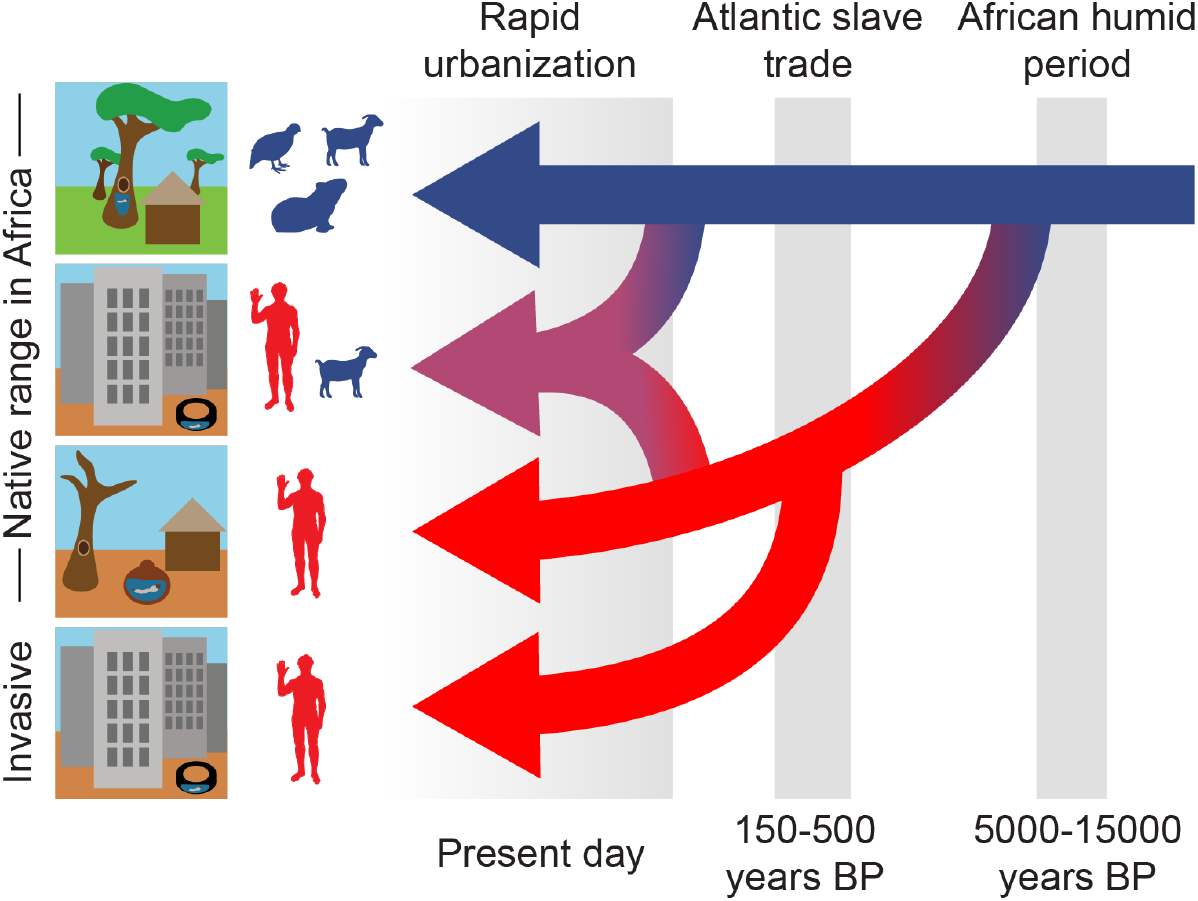
Three epochs define the origin and spread of the human-specialist form of *Ae. aegypti*. Our analyses suggest that the human-specialized *Ae. aegypti* rapidly diverged from their generalist counterparts after the end of the African Humid Period, with the emergence of settled human societies in intensely seasonal habitats with long, hot dry seasons. Human specialists then migrated from West Africa to the Americas during the Atlantic Slave Trade. Finally, we find evidence of a recent influx of human-specialist ancestry into the rapidly growing cities consistent with an ongoing shift in the ecology of *Ae. aegypti* in present-day Africa.

The finding that human-specialists emerged at the end of the African Humid Period is consistent with the hypothesis that climate provided the initial selective push towards specialization. It was at this time, approximately 5,000 years ago, that the Sahara dried and the highly seasonal Sahel biome to its south came to be. *Ae. aegypti* mosquitoes inhabiting the area would have found themselves in a novel situation, devoid of rain-filled breeding sites for nine months of the year. This could have driven adaptation to human-derived breeding sites (and also perhaps human-derived blood meals) in order to survive. In addition to the signal of rapid divergence starting about 5,000 years ago, we saw some evidence for steady population growth and accumulation of modest genetic structure starting 15,000 to 30,000 years ago. This observation is compatible with a model where some populations expanded into newly habitable parts of the present day Sahara/Sahel at the start of the African Humid Period, giving rise to weak geographical structure, before rapidly diverging in response to a drying climate and the emergence of the human-specialist niche at its end. Although they seem to have occasionally interbred, specialist and generalist populations then became largely isolated and ecologically distinct.

It is widely accepted that human-specialist *Ae. aegypti* spread to the Americas (and eventually throughout the global tropics) during the Atlantic Slave Trade, ushering in a new era of vector-borne disease (*6*). We relied on this historical understanding to calibrate our coalescent scaling factor. However, our results also help establish its credibility. The cross-coalescent signal between Ngoye, Senegal and Santarem, Brazil closely fit *a priori* expectations, even without calibration. Using literature values for average insect mutation rates and our best estimate of *Ae. aegypti* generation time would have also yielded good correspondence between the inferred timing of migration and the historical record of the Atlantic Slave Trade (Figure 2A). The duration and sudden drop-off in migration observed in the coalescent signal also closely matches expectations from the historical record (Figure 2A).

While our cross-coalescent analyses are remarkably consistent with previous expectations, several caveats and open questions remain. For example, imagine the main migration pulse of *Ae. aegypti* out of Africa was actually more recent than the Atlantic Slave Trade. In this case, divergence between human-specialists and generalists would have also been more recent than we infer here, perhaps corresponding to some event after the end of the African Humid Period such as expanding human populations. Breeding in human-stored water may have been forced by climate, but it also requires the presence of settled human societies. Interestingly, our 5,000 year estimate also corresponds closely to the estimated timing of pearl millet domestication in the western Sahel, an event that was likely also precipitated by changes in climate at that time (*36*). Multiple aspects of the current human-specialist niche thus came together simultaneously in this region.

Populations with substantial human-specialist ancestry have also been described in Luanda, Angola, which was a major port during the Atlantic Slave Trade (*6, 29*). These populations were likely the proximate source of many *Ae. aegypti* introduced to the Americas (*29*), raising the question of whether human-specialist populations of *Ae. aegypti* could have arisen outside the Sahel. However, the closest generalist relatives of human-specialist populations are found in West Africa (*29*), and there was no analogous sudden emergence of a human-specialist niche around Angola 5,000 years ago. Although present distributions of populations may not perfectly match their historical distributions, the most parsimonious explanation is that human-specialist populations (the “proto-*Aedes aegypti aegypti”* populations described in ref. 5) first arose about 5,000 years ago in the West African Sahel and later spread to both Luanda and the Americas in association with the Atlantic Slave Trade. The timeline of divergence of human-specialist *Ae. aegypti* could be further clarified by direct estimation of the *de novo* mutation rate and better ecological studies of generation times in the field. Finally, while the analyses presented here are based on patterns of neutral drift within and between populations, future work on the genetic basis of traits that adapt *Ae. aegypti* to human hosts and habitats could shed light on the timing of key selective events.

This study represents one of the first applications of cross-coalescent analysis to a non-model system. Cross-coalescent analyses are a powerful tool for understanding historical relationships between populations, but they are very difficult to implement in practice, largely due to the logistical/computational burden of developing a large genome phasing panel and the difficulty of estimating an accurate coalescent scaling factor. The correct calibration of coalescent analysis is notoriously difficult even in well-understood systems like humans (*37*). Here, we take advantage of a continent-wide set of genomes, combined with read-based prephasing and population-wide statistical phasing to develop a phasing panel that can enable future studies in *Ae. aegypti* with a lower computational and logistical barrier to entry. Furthermore, our approach to coalescent scaling factor calibration based on the historical record could likely be applied in other taxa whose history is characterized by known historical events as well as events with unknown timing. Similar calibrations have been used to understand the relationship between coalescent analyses and human history (*38, 39*), but this approach may prove especially valuable in non-model organisms where population genomic parameters are less well characterized.

Human-specialization first emerged within *Ae. aegypti* ∼5,000 years ago, but now, in the face of rapid urbanization of sub-Saharan Africa, generalist populations of this species seem to be undergoing a new shift. We estimate an influx of human-specialist alleles into the rapidly growing cities of Kumasi and Ouagadougou within the past 20-40 years. Uncertainty remains about the actual geographical origin of the human-specialist ancestry tracts found in both cities. They are likely derived from local gene flow from nearby specialist populations, with whom both KUM and OGD showed affinity in previous *ADMIXTURE* analyses (*2*). However, they could alternatively (or additionally) be derived from distant specialist populations or even invasive populations outside Africa; dormant *Ae. aegypti* eggs are often transported inadvertently, e.g. in the used-tire trade. More comprehensive sampling and analysis of human-specialist populations inside and outside Africa will be necessary to identify the precise origins of the genetic material currently transforming urban populations in Africa.

Regardless of the source, the influx of human-specialist ancestry into *Ae. aegypti* mosquitoes inhabiting African cities will likely have important implications for human health. Human-specialist *Ae. aegypti* are both more likely to bite humans and more likely to serve as competent vectors of dengue, Zika, and yellow fever (*2, 40*–*42*). Ouagadougou has seen repeated outbreaks of dengue fever in recent years (*43, 44*), and mosquitoes there are still genetically and phenotypically intermediate between generalists and specialists (*2, 29, 44*). It is possible that such populations will eventually become more specialized as urbanization continues. Likewise, the emergence of urban yellow fever since 2016 in Africa may also have been hastened by the expansion of human-specialist populations of *Ae. aegypti* (*45, 46*). For these reasons, monitoring and control of *Ae. aegypti*, taking into account possible changes in the vector capacity of local populations over time, should remain a high priority in the native range of this key vector species.

## Methods

### Genome phasing

We developed a genomic phasing panel using 389 previously published *Ae. aegypti* genomes sequenced to 15x coverage (*2*), including 359 from across the species’ native range in Africa, as well as 30 from invasive populations outside of Africa (16 from Bangkok, Thailand and 14 from Santarem, Brazil) (*2*). This panel also included 4 *A. mascarensis* individuals, but phasing quality for these individuals may be lower due to the small number of individuals used. We used as input the full set of previously *bcftools*-called biallelic SNPs (*2*). We further filtered the panel to include variant sites with GQ>20 and DP>8, including only variants in putative single-copy regions (mean coverage 5-30x), and excluding annotated centromeric and repeat regions, as well as the sex locus (*35*). Ultimately, our phasing panel comprised 52,534,217 biallelic SNPs. We used a two-step phasing procedure. We first pre-phased nearby heterozygous sites using information present in sequencing reads within individuals using *HAPCUT2* (*20*), and then carried out statistical phasing on a population level with *SHAPEIT4*.*2* (*23*), using a phase set error rate of 0.0001. Due to memory constraints, we carried out two separate *SHAPEIT4*.*2* runs – one with all samples from East and Central Africa, and a second with all samples from West Africa and outside of Africa. This strategy should maximize the use of phasing information from samples showing similar patterns of linkage disequilibrium (*47*).

### MSMC2 and MSMC-IM analyses

We used MSMC2 and MSMC-IM according to published best practices to carry out cross-coalescent and isolation-with-migration analyses, respectively (*21, 48*). For this family of analyses, inference may be less accurate if the recombination rate is much higher than the mutation rate. This constraint is not a problem for the human genomic data that these analyses were originally designed for, but may present a barrier to analyses in other species (*49*). For *Ae. aegypti*, both recombination and mutation rates are likely about an order of magnitude lower than those observed in humans (*15, 35*), suggesting a similar ratio to that observed in humans (∼1) and providing support for the application of these analyses to *Ae. aegypti*. We first generated genome-specific masks for each genome using the script bamCaller.py, which is provided with the MSMC2 package. We also used a general genomic mask that included only putative single-copy regions (mean coverage 5-30x) and excluded annotated centromeric and repetitive regions. This general genomic mask further excluded regions previously identified as under selection between specialists and generalists (*2*), as well as the sex locus, and sites that were annotated as uncallable using SNPable on the African consensus of the AaegL5 assembly, with parameters -l 150 -r 0.5 (*50*). We used bcftools to extract phased genomes for focal individuals from our phasing panel. We then carried out cross-coalescent analyses using MSMC2. We carried forward the resulting MSMC2 output into MSMC-IM analysis, fitting isolation-with-migration models to the fitted cross-coalescent estimates, with default regularization settings. We bootstrapped MSMC2 and MSMC-IM analyses using ten replicates of three 400 Mb chromosomes composed of resampled blocks of 20 Mb.

We characterized rates of cross-coalescence and historical effective population sizes among five populations. We used Ngoye, Senegal (NGO) as a representative of human-specialists living in Africa. We used Ouahigouya, Burkina Faso (OHI) and PK10, Senegal (PKT, a forest area outside of Kedougou, Senegal) as representatives of nearby generalist populations in West Africa. We used Bangkok, Thailand (BKK) and Santarem, Brazil (SAN) as representatives of invasive human-specialist populations outside of Africa. For each population, we used two representative individuals, one male and one female. We confirmed the sex of these individuals by checking rates of read mapping at the sex locus (chromosome 1, positions 151680000-152950000) — males all had >700 counts per million, and females all had <400 counts per million. Note, however, that the sex locus was masked for coalescent analyses.

As discussed in the Results, our cross-coalescent analyses yielded results in coalescent units that need to be scaled by mutation rate and generation time to yield dates for key events.These parameters are difficult to calibrate even in well-characterized systems like humans and *Drosophila*. For this reason, we used the well-described relationship between the Atlantic slave trade and the spread of *Ae. aegypti* out of Africa to ground-truth and calibrate our coalescent scaling factor. We used the Bhattacharyya coefficient (implemented in a custom R function) to measure the degree of concordance between the inferred *Ae. aegypti* migration rate between West Africa and the Americas (using Ngoye, Senegal and Santarem, Brazil as reference populations) and the intensity of the Atlantic slave trade as documented in the Slave Voyages database (*30*). We then used the coalescent scaling factor that gave the strongest concordance between inferred migration and slave trade intensity (corresponding to a mutation rate of 4.85×10^−9^, assuming 15 generations per year) to calibrate our subsequent analysis of divergence between human-specialist and generalist lineages.

The climate history of the Sahara and Sahel regions has been reconstructed from analysis of plant-derived marine sediments from the Eastern Atlantic (*31*). We visualized the African Humid Period by using *ImageJ/FIJI* (*51*) to trace the four reconstructed records of precipitation over the past 25,000 years (GC27, GC37, GC49, GC69) in the Sahel and Sahara described by Tierney et al. (*31*). We averaged these traces, interpolating levels of precipitation with a 100-year step using the *R* function *approxfun* (*52*), and plotted the rolling average of this trace with a 1000-year window and 100-year step.

### f3 analyses

In order to determine how best to model potentially admixed urban populations in KUM and OGD, we used *f3* analyses of our previously called set of genome-wide SNPs with minor allele frequency >0.05 (*2*), as implemented in the program *threepop* (*53*), to test for signatures of admixture between different human-specialist (BKK, SAN, NGO) and generalist (OHI, KIN) populations. We used block-jackknifed Z-scores to assess whether the resulting values were significantly negative (*32*).

### AncestryHMM analyses

We used AncestryHMM to detect tracts of human-specialist ancestry in KUM and OGD, using a single-pulse model, running a separate analysis for each chromosome. We filtered the previously described biallelic SNP set (*2*) to include only ancestry-informative sites characterized by an allele-frequency difference between populations of >0.3 and a minimum of 10 sampled alleles per population. Sites were then thinned such that no two were closer than 10,000 bp apart,. We estimated recombination rates between sites using the recombination map published with the AaegL5 genome assembly (*35*). For each analysis, we used reference panels of 10 unrelated and unadmixed OHI individuals and 16 unrelated and unadmixed BKK individuals (*2*). *AncestryHMM* requires estimates of admixture proportions for each chromosome as part of its input. We estimated admixture proportions on each chromosome using the set of 1,000,000 unlinked variants used to estimate genome-wide admixture proportions in a previous *ADMIXTURE* analysis (*2, 54*), subsetting these variants for each chromosome. *AncestryHMM* simultaneously estimates admixture timing during model training – we used 80 bootstraps resampling blocks of 1000 ancestry-informative sites to construct 95% confidence intervals for admixture pulse timing for each chromosome in each population. Since selection associated with adaptation to urban habitats could shape lengths of admixture tracts, we masked regions previously identified as under selection between human-specialists and generalists when estimating admixture timing.

### Comparison of admixture patterns between cities

In order to determine whether the distribution of human-specialist ancestry across the genome was more similar between OGD and KUM than would be expected from random chance, we used circular chromosome permutation tests (*55*). We first used Viterbi posteriors from each individual within a single population to calculate a population-specific local ancestry proportion at each site used in our *AncestryHMM* analysis, and then calculated the Pearson correlation of local ancestry in KUM with that in OGD across all genomic loci. In each permutation, we randomly selected a new starting point for each chromosome, placing the preceding sequence at the end of the permuted chromosome but otherwise leaving the spatial relationship between loci undisturbed. We then recorded the Pearson correlation coefficient between the permuted distributions of ancestry in KUM and OGD, constructing a null distribution across 1000 permutations.

### AncestryHMM simulations

In order to be sure that the distribution of admixture tracts found by our *AncestryHMM* analysis reflects actual differences in local ancestry, and not differences in our power to detect them, we carried out a series of simulations taking advantage of our phasing panel. We simulated heterozygous tracts of different sizes (500kb, 1Mb, 2Mb, 10Mb) derived from the first haplotype of a phased BKK genome (sample Debug_010_aegypti_Bangkok_Thailand_01) on the background of two different unadmixed OHI individuals (Debug023_aegypti_Ouahigouya_BurkinaFaso_F_11, Debug023_aegypti_Ouahigouya_BurkinaFaso_M_10), using *bedtools random* to generate random intervals in which to place the tracts, *bcftools* to extract the tracts from the BKK genome, and *bedtools intersect* to generate simulated input files for *AncestryHMM*. For each of the four tract lengths, we carried out 100 replicate simulations. We noticed that *AncestryHMM* found tracts of specialist ancestry across all simulations using a given OHI individual in some small regions of the genome. However, these regions differed between the two OHI individuals used in our simulations. We therefore suspect these regions are small *bona fide* tracts of human-specialist ancestry in these otherwise unadmixed OHI individuals rather than general features of the analysis or reference panels used.

## Data availability

Scripts and processed data are available at github.com/noahrose/aaeg-evol-hist. Raw genomic data are available in the NCBI SRA at accession PRJNA602495.

---

**Figure S1.**
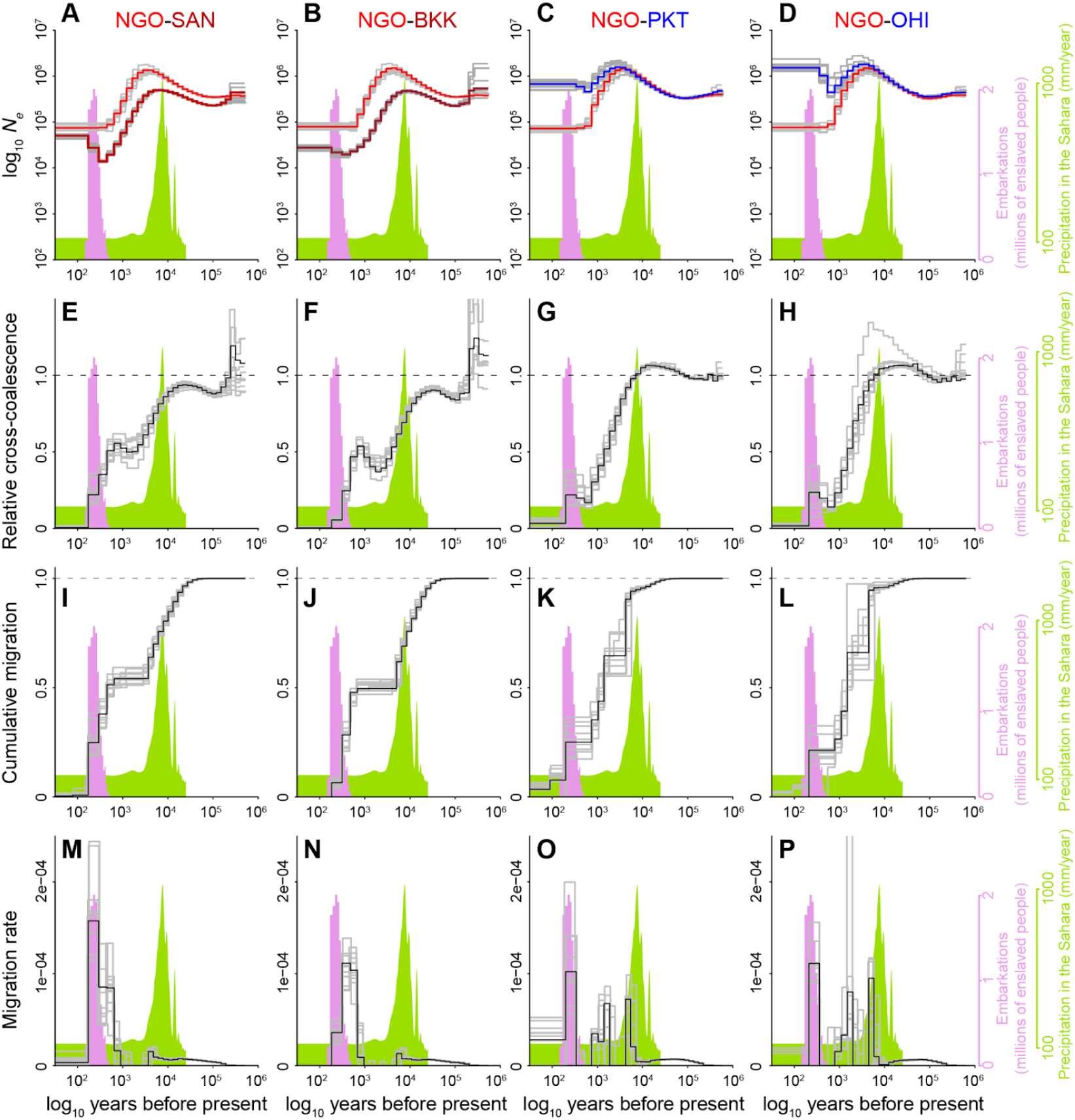
Full detail of cross-coalescent analyses comparing the Sahelian human-specialists to either invasive human specialists or nearby generalists. For each population pair (i.e. column), we show MSMC2-inferred changes in effective population size over time (**A**–**D**), MSMC2-inferred relative cross-coalescence (expected to plateau at 1 when populations have merged into a single ancestral population) (**E**–**H**), MSMC-IM-inferred cumulative migration (also expected to plateau at 1 when populations have merged into a single ancestral population) (**I**–**L**), and MSMC-IM-inferred migration rate (fraction per generation) (**M**–**P**). First two columns show results for the Sahelian human specialist population NGO compared to invasive human-specialist populations from Brazil (SAN; panels A, E, I, M) or Thailand (BKK; panels B, F, J, N). Note that the invasive populations have both recent (hundreds of years, as expected) and much older (thousands of years) pulses of coalescence with NGO. The older signals likely arise from the fact that both NGO and invasives have somewhat mixed ancestry. NGO has experienced substantial admixture from nearby generalists in recent years and is therefore an imperfect proxy for the African human-specialists that gave rise to invasive populations. Likewise, invasive populations have likely experienced gene flow from other African generalist populations (e.g. in Angola, a major slave trading hub) *(23)* that diverged from West African generalists long before they gave rise to specialists. The last two columns show results for the Sahelian human specialist population NGO compared to nearby generalist populations from Senegal (PKT; panels C, G, K, O) or Burkina Faso (OHI; panels D, H, L, P). In addition to the signal of rapid divergence/migration ∼5,000 years ago, we also observe increased divergence/migration during the Atlantic Slave Trade – this signal may reflect increased movement within the continent also associated with the Atlantic Slave Trade, or could reflect other changes in patterns of gene flow during this period of time.

**Figure S2.**
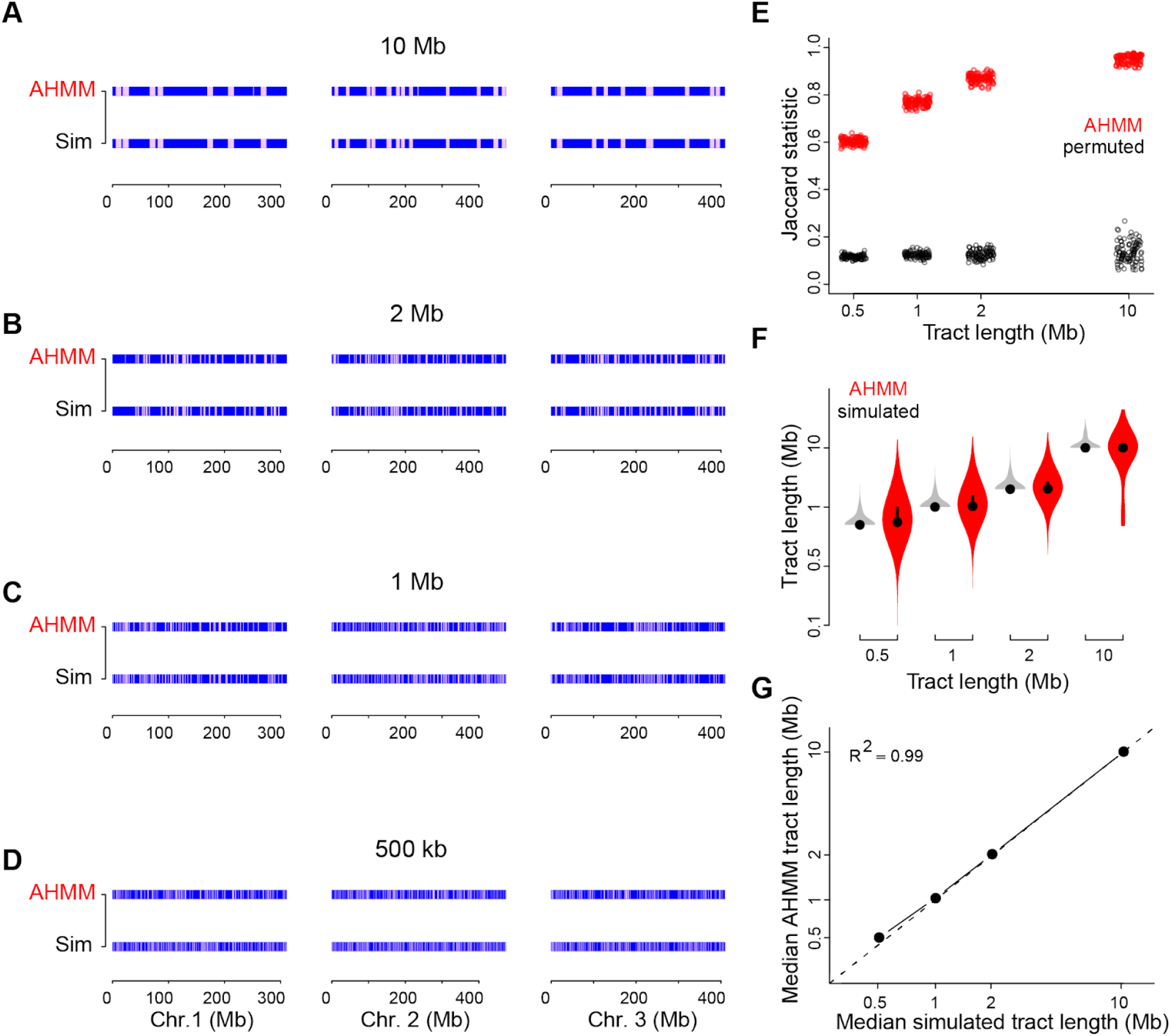
Simulations confirm the reliability of admixture analyses in West African cities. **A**–**D**, AncestryHMM (AHMM) analysis (top row of each panel) reliably detected simulated tracts of BKK ancestry (bottom row of each panel) in a OHI background with tract lengths of 500 kb (**A**), 1 Mb (**B**), 2 Mb (**C**), and 10 Mb (**D**). **E**, Overlap between simulated and AHMM-inferred tracts, as measured by the Jaccard statistic, which should take a value of zero in the case of no overlap and one in the case of perfect overlap. Overlap was strong for all four tract lengths (all permutation P<0.01), though longer tracts were more accurately detected. **F**, AHMM-inferred tracts show a similar mean length to simulated tracts, though AHMM showed a broader distribution of tract lengths overall. Simulated tracts (grey) were all the same length, but overlapping tracts were merged, leading to some longer tracts. **G**, Median simulated and AHMM-inferred tract lengths are concordant.

**Figure S3.**
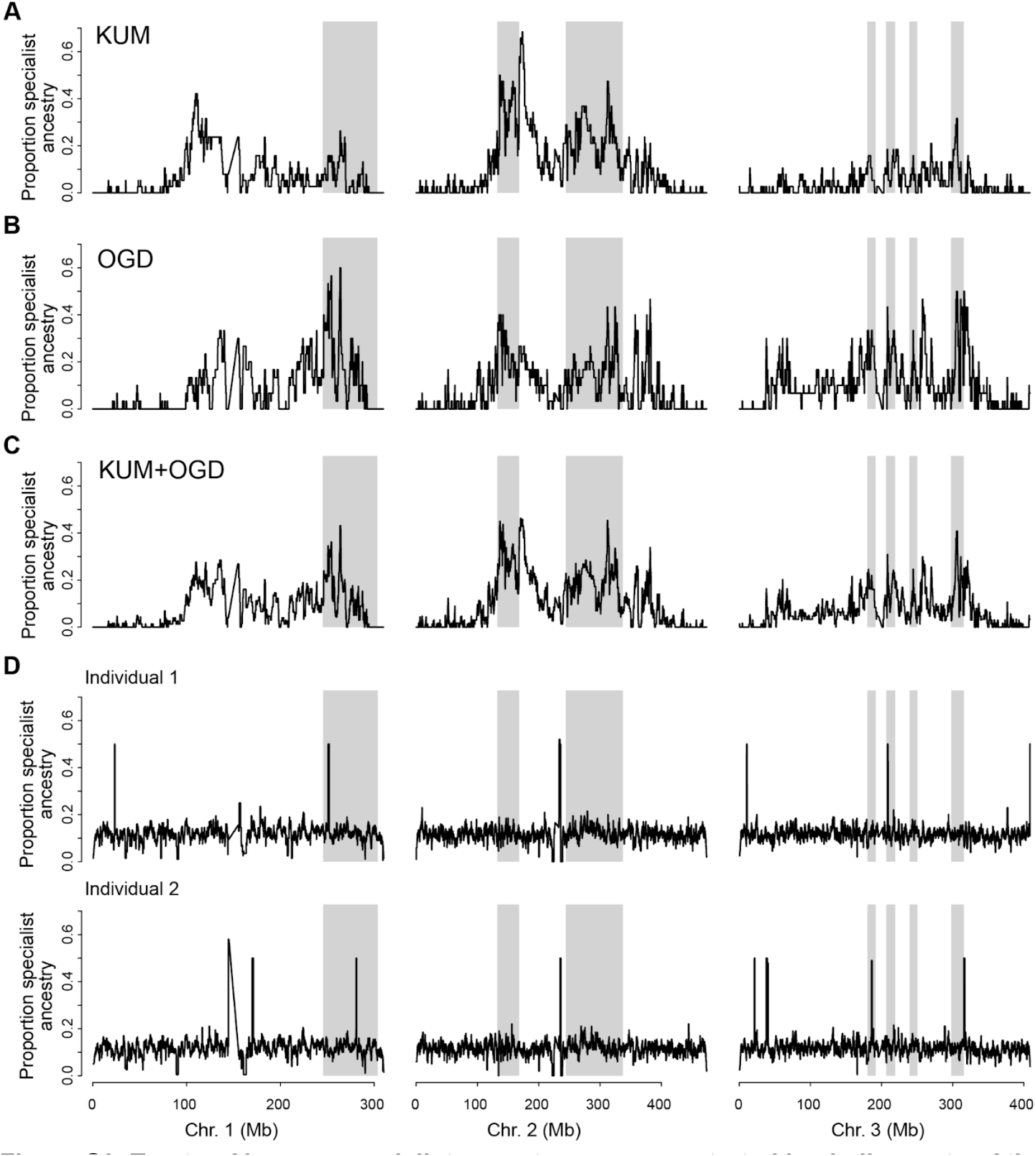
Tracts of human-specialist ancestry are concentrated in similar parts of the genome in Kumasi and Ouagadougou. **A**–**B**. Local fraction of ancestry derived from human specialists across the genome in KUM (**A**) and OGD (**B**) (using BKK and OHI as specialist and generalist reference panels, respectively). **C**. As in panel A, but averaging across KUM and OGD. **D**. Randomly simulated tracts of human specialist ancestry (160 tracts of 2Mb length per simulation, similar to the conditions observed in KUM and OGD) are evenly detected across the genome of two OHI individuals. A few small heterozygous tracts are detected across all simulations regardless of whether simulated tracts were placed in these locations, likely representing rare human-specialist tracts present in this otherwise unadmixed individual. The tracts detected across all simulations are in different locations in the two different individuals, providing further evidence that these signals are caused by rare human-specialist tracts in OHI and do not reflect a systematic feature of the analysis or reference panels. Grey shading represents the outlier regions previously identified as involved in human specialization (*2*).

## Notes

### Competing Interest Statement

The authors have declared no competing interest.

